# Differentiation of human induced pluripotent or embryonic stem cells decreases the DNA damage response

**DOI:** 10.1101/094938

**Authors:** Kalpana Mujoo, Raj K. Pandita, Anjana Tiwari, Vijay Charaka, Sharmistha Chakraborty, Walter N. Hittelman, Nobuo Horikoshi, Clayton R. Hunt, Kum Kum Khanna, Alexander Y. Kots, E. Brian Butler, Ferid Murad, Tej K. Pandita

## Abstract

Genomic integrity is critical for preservation of stem cell function and is, maintained through a robust DNA damage response (DDR) with systemized DNA DSB repair by either the non-homologous end joining (NHEJ) pathway or the homologous recombination (HR) pathway. To examine DDR during stem cell differentiation, human embryonic (hES) and induced pluripotent stem (IPS) cells were exposed to DNA damaging agents and DNA damage signaling/repair measured. Differentiated cells displayed a higher frequency of residual DNA damage, chromosomal aberrations, cells with delayed γ-H2AX foci disappearance and a reduced number of RAD51 foci. Factors impacting DNA DSB repair by HR formed reduced foci in differentiated cells. The reduction in repairosome foci formation after DNA damage was not due to changes in HR protein levels, which were unchanged by differentiation. Differentiated cells also displayed a higher frequency of stalled DNA replication forks and decreased firing of new replication origins from transient inhibition of DNA synthesis by hydroxyurea treatment. In addition, we observed that differentiated cells exhibit a higher frequency of R-loops. A similar decline in DDR was observed as early stage mouse astrocytes differentiated into later stage astrocytes. Our studies thus suggest that DSB repair by homologous recombination is increasingly impaired during stem cell differentiation while the NHEJ pathway is minimally altered.

## Introduction

Stem cells have the dual ability to self-renew over the lifetime of an organism and also to differentiate into multiple cell lineages (Weissman, Anderson, and Gage 2001, Seita and Weissman 2010). The majority of mammalian cells *in situ* originate from a corresponding progenitor cell but are terminally differentiated. Various factors, including reactive oxygen species (ROS), which accumulate during differentiation and over the stem cell lifespan, can cause DNA damage (Mikhed et al. 2015). In addition, differentiation-dependent changes in chromatin structure and transcription (Nashun, Hill, and Hajkova 2015, Tran et al. 2015); can also impact genomic integrity by altering the DNA damage response (DDR) and repair. Thus, genomic stability is likely to be under increased stress during differentiation. There is no information available whether differentiation of stem cells impacts DSB repair and we studied in detail the mechanisms by which pluripotent stem cells versus differentiated cells respond to double strand breaks (DSB) induced by DNA damaging agents in detail in isogenic differentiating cell lines.

Stem cells benefit throughout their lifetime from a robust DNA damage repair activity that enhances resilience towards various environmental factors. Indeed, somatic cells and stem cells differ in their radio-sensitivity (Chlon et al. 2016, Maynard et al. 2008, Lan et al. 2012, Momcilovic et al. 2009, Wilson et al. 2010), however, it is not known whether DNA DSB repair is impacted during the stem cell differentiation. In order to understand the relationship between stem cell differentiation and DNA damage repair, we compared DNA damage responses and DNA repair pathways in human embryonic (H9) and induced pluripotent stem cells (B12-2 and B12-3) with their isogenic differentiated progeny and found that DNA damage repair by HR is significantly reduced in differentiated cells.

## Results

### Characterization of differentiation markers in iPS cells

We used human iPS cell lines B12-2 and B12-3 and ES cells to compare the DDR between undifferentiated and differentiated cell status. The cell lines used were positive for OCT4 or Nanog (Fig. 1A) and cell markers (ectoderm β-III tubulin, TUJ1; mesoderm smooth muscle actin, SMA; and endoderm alpha-feto protein, AFP) confirmed embryoid body (EB) directed differentiation into the three germ layers (Fig. 1B). Western blot analysis revealed a time-dependent decrease in OCT4 and nanog (Fig. 1C) as well as hMOF protein levels during differentiation (Fig. 1D). hMOF acetylates histone H4 lysine 16, levels of which were also reduced in differentiated cells (Fig. 1D), and plays a role in regulating stem cell differentiation as well as in DNA double strand break repair (Gupta et al. 2008, Kumar et al. 2011, Thomas et al. 2008, Li et al. 2012).

**Figure 1:**
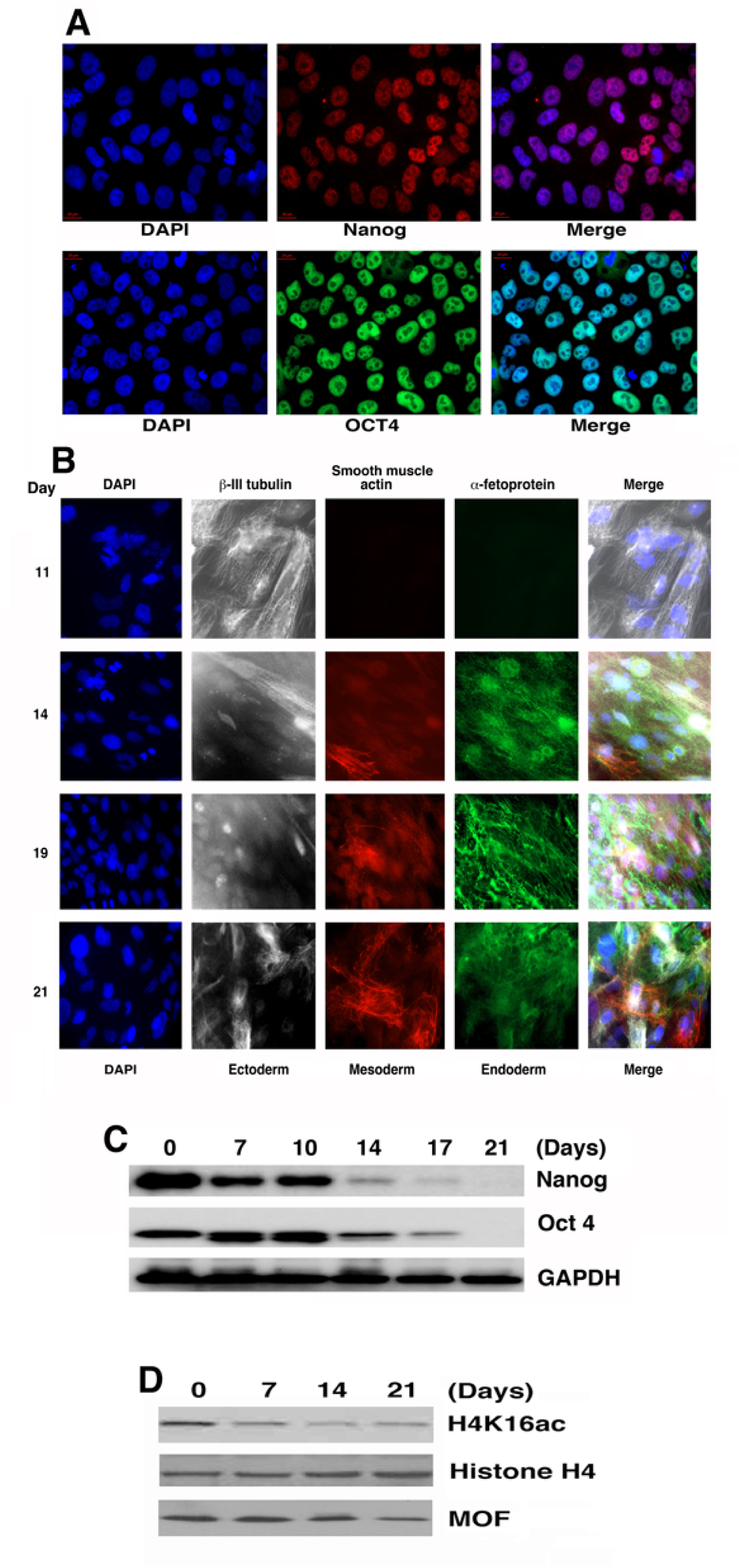
Stem cell markers and their differentiation. **(A)** Immunostaining with antibody against Nanog and OCT4 in iPS cells. **(B)** Immunostaining with different antibodies to detect stem cell differentiation into three germ layers. **(C)** Western blot showing Nanog and Oct 4 levels during various stages of differentiation and; **(D)** Western blot showing MOF, Histone H4 and H4K16ac levels during temporal differentiation.

### DNA DSBs in undifferentiated and differentiated stem cells

iPS cells were exposed to graded doses of IR and DSBs were measured by the comet assay under neutral conditions. An identical linear relationship between the comet tail moment and radiation dose was observed in undifferentiated and differentiated cells (Fig. S1), indicating similar DSB induction. Cells were then examined for ability to rejoin the DSBs post-irradiation. A significant increase in residual DSBs was observed in differentiated cells (Fig. 2 A and B), suggesting that the cells have a reduced DSB repair capacity in comparison to undifferentiated cells.

**Figure 2:**
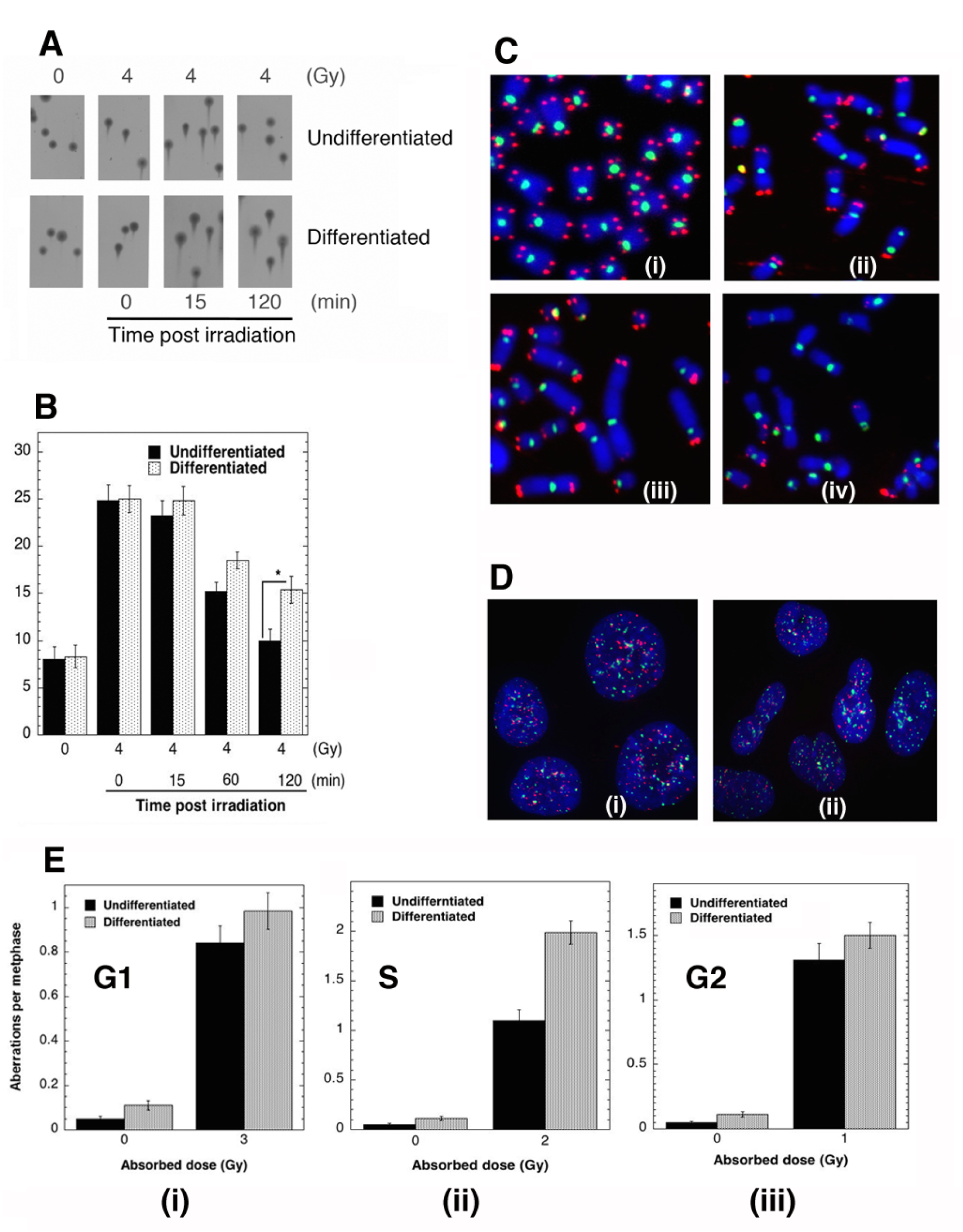
Detection of chromosome and DNA damage in stem cells and differentiated cells. DSBs in stem and differentiated cells after exposure to 4 Gy were detected by comet assay at intervals up to 120 min **(A)**. **(B)** Quantitative analysis of mean tail moment (time course) in stem cells and differentiated cells post exposure to IR. Error bars indicate ± SEM. Significance using paired student t-test is shown. * *P*< 0.05. **(C)** Micronuclei, chromosome blebbing, aneuploidy and polyploidy **(D)** metaphase chromosomes-telomere and centromere FISH in (i) undifferentiated stem cells; (ii) differentiated cells showing loss of telomeres and dicentric type chromosome aberrations; (iii) fusion of chromosomes and dicentric; (iv) loss of telomeres and polyploidy. **(E)** Analysis of cell cycle specific residual chromosome damage in undifferentiated and differentiated cells with chromosomal aberration analyzed at metaphase; G1, S, and G2 cell cycle phase aberrations include dicentrics, centric rings, interstitial deletions/acentric rings, and terminal deletions; all categories of asymmetric chromosome aberrations were scored; differentiated cells showed significantly higher chromosomal aberration frequencies compared to control cells (\*\**P* > 0.01, Student t test, n=3).

### Chromosome aberration analysis

We determined whether the increased residual DSBs observed in differentiated cells correlated with chromosome aberrations by measuring basal level and IR-induced chromosome aberrations at metaphase. Differentiated cells were found to have a significantly higher frequency of chromosome aberrations (breaks, gaps, radials, dicentrics, aneuploids and polyploids) as compared to undifferentiated cell (Fig. 2C and Fig. S2B). About 8-12 % of differentiated cells had chromatin blebbing (Fig. 2D) and endo-reduplicated chromosomes (Fig. S2A). We determined whether the increased frequency of aberrations seen in differentiated cells is due to a defect in repair in a specific DSB repair pathway by comparing the cell cycle specificity of IR-induced chromosome aberrations. No significant differences were observed between undifferentiated and differentiated cells in IR-induced G1-specific aberrations (Fig. 2Ei), suggesting the NHEJ repair pathway is largely intact. However, the frequency of S-phase specific chromosome aberrations was significantly higher in differentiated as compared to undifferentiated cells (Fig. 2Eii) while G2-type chromosomal aberrations did not show any significant differences (Fig. 2Eiii). Since DSB repair by HR is active in S phase, the present data suggests that differentiated cells may have defects in the HR repair pathway.

### DDR in stem cells before and after differentiation

The initial events of DDR are ATM autophosphorylation and H2AX phosphorylation at Ser139 (*γ*-H2AX), foci of the latter often serving as a maker for the presence of a DSB. The frequency of IR-induced γ-H2AX foci induction was identical in undifferentiated and differentiated cells; however, there was a prolonged delay in the disappearance of *γ*-H2AX foci in differentiated cells (Fig. 3A), indicating compromised repair. The induction of IR-induced *γ*-H2AX foci was identical in differentiated ES (Fig. S3 A) and iPS cells (Fig. 3A), suggesting the overall sensing of DNA damage is unaltered in differentiated cells. The higher frequency of residual *γ*-H2AX foci in differentiated cells, however, indicates a defect in DSB repair that could lie in either the non-homologous end joining (NHEJ) and/or HR pathway.

**Figure 3:**
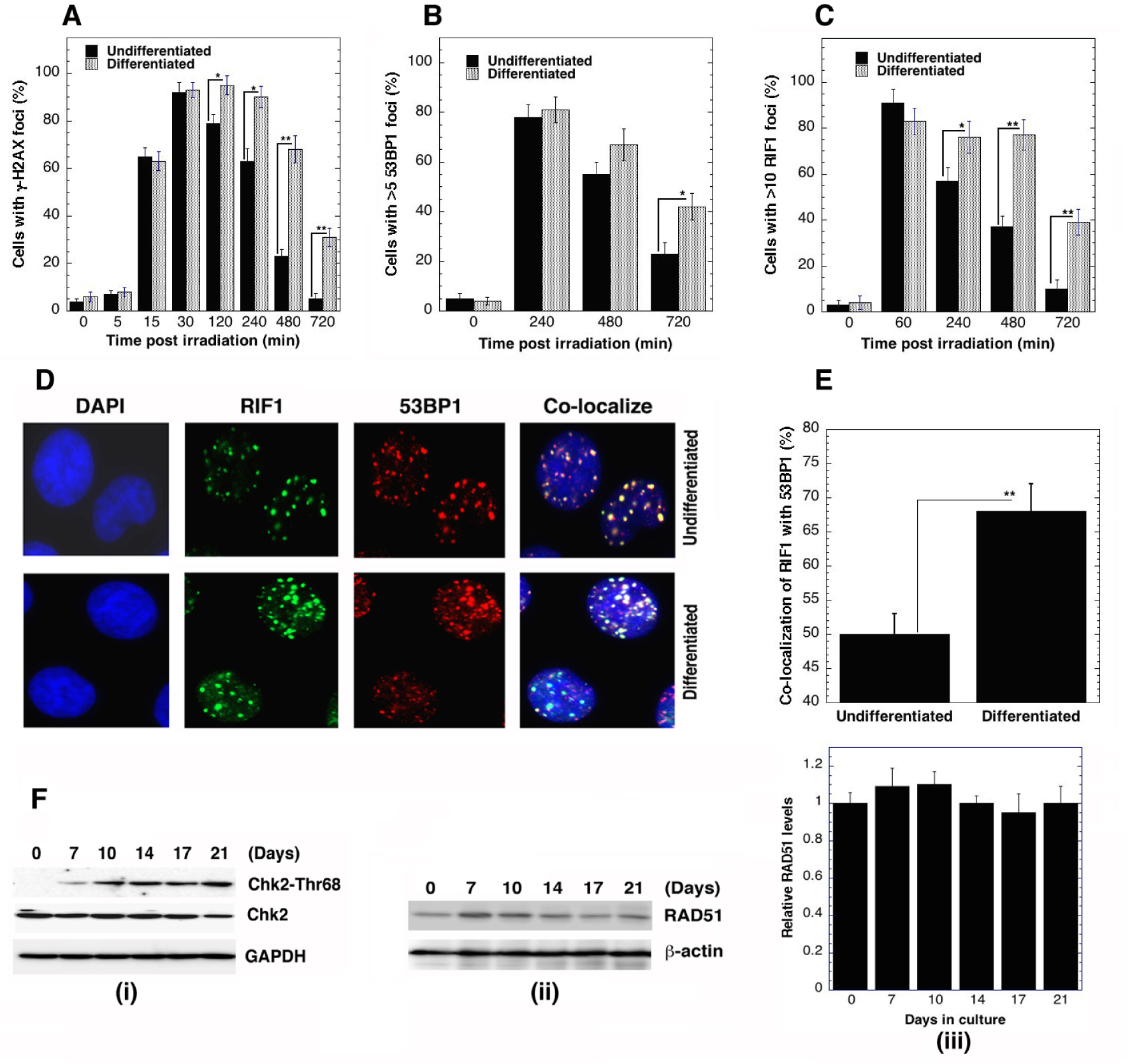
DNA damage response in iPSCs and differentiated cells. B12-2 and differentiated cells were exposed to various doses of IR **(A)** γ-H2AX (2 Gy), **(B)** 53BP1 (6 Gy), and **(C)** RIF1 (6 Gy), then stained and analyzed for foci formation. **(D)** Co-localization of 53BP1 and RIF in B12-2 and differentiated cells. **(E)** Quantitation of RIF1 and 53BP1 co-localization; 53BP1 and RIF1 foci were counted for 3 sets of 25 cells and percentage of co-localized 53BP1/RIF1 foci were calculated relative to total number of foci (53BP1+RIF1); western blotting of Chk2 **(Fi)**, RAD 51 **(Fii)** and histogram showing relative levels of RAD51 (mean of three blots) **(Fiii)** during differentiation of iPS cells.

### Repairosome foci analysis of HR-related protein factors

The 53BP1 protein has been implicated in the suppression of HR (Morales et al. 2003, Ward et al. 2003, Zimmermann et al. 2013), and the first downstream effector of 53BP1 is RIF1 (Chapman et al. 2013, Di Virgilio et al. 2013, Escribano-Diaz and Durocher 2013, Escribano-Diaz et al. 2013, Feng et al. 2013, Zimmermann et al. 2013). We observed that while the initial formation of 53BP1 foci post-irradiation was identical, there was a significant delay in 53BP1 foci clearance in differentiated cells (Fig. 3B). The frequency of RIF1/53BP1 foci colocalization was also higher in differentiated cells as compared to undifferentiated cells (Fig. 3 C-E). Accumulation of RIF1 at DSB sites containing phosphorylated 53BP1 (Anbalagan et al. 2011, Bonetti et al. 2010) inhibits the DNA resection step of HR (Bunting et al. 2010), suggesting differentiation-dependent suppression of DSB repair by HR. Cellular levels of RAD51 and Chk2 proteins are similar in differentiated versus undifferentiated cells but Chk2 phosphorylation at Thr68 increases during differentiation (Fig. 3Fi). Despite the identical RAD51 levels (Fig. 3Fiii), there was a significant decrease in RAD51 foci formation after irradiation in differentiated ES cells (Fig. S3 B). Consistent with the reduced frequency of cells with IR-induced RAD51 foci, we also observed reduced IR-induced BRCA1 MRE11, RAP80, and FANCD2 foci in differentiated cells (Fig. 4 A-F), supporting the argument that differentiated stem cells have a reduced ability to repair DSBs by HR. We further examined whether the DDR is similarly altered when primitive cells derived from a developing animal organ are fully differentiated *in vitro*. Later stage astrocytes had a higher frequency of cells with delayed disappearance of *γ*-H2AX foci and a reduced number of RAD51 foci indicating that differentiated astrocytes showed a similar DDR defect as observed in the stem cell-derived differentiated cells (Fig. S4), suggesting decreased HR is a general feature of cell differentiation.

**Figure 4:**
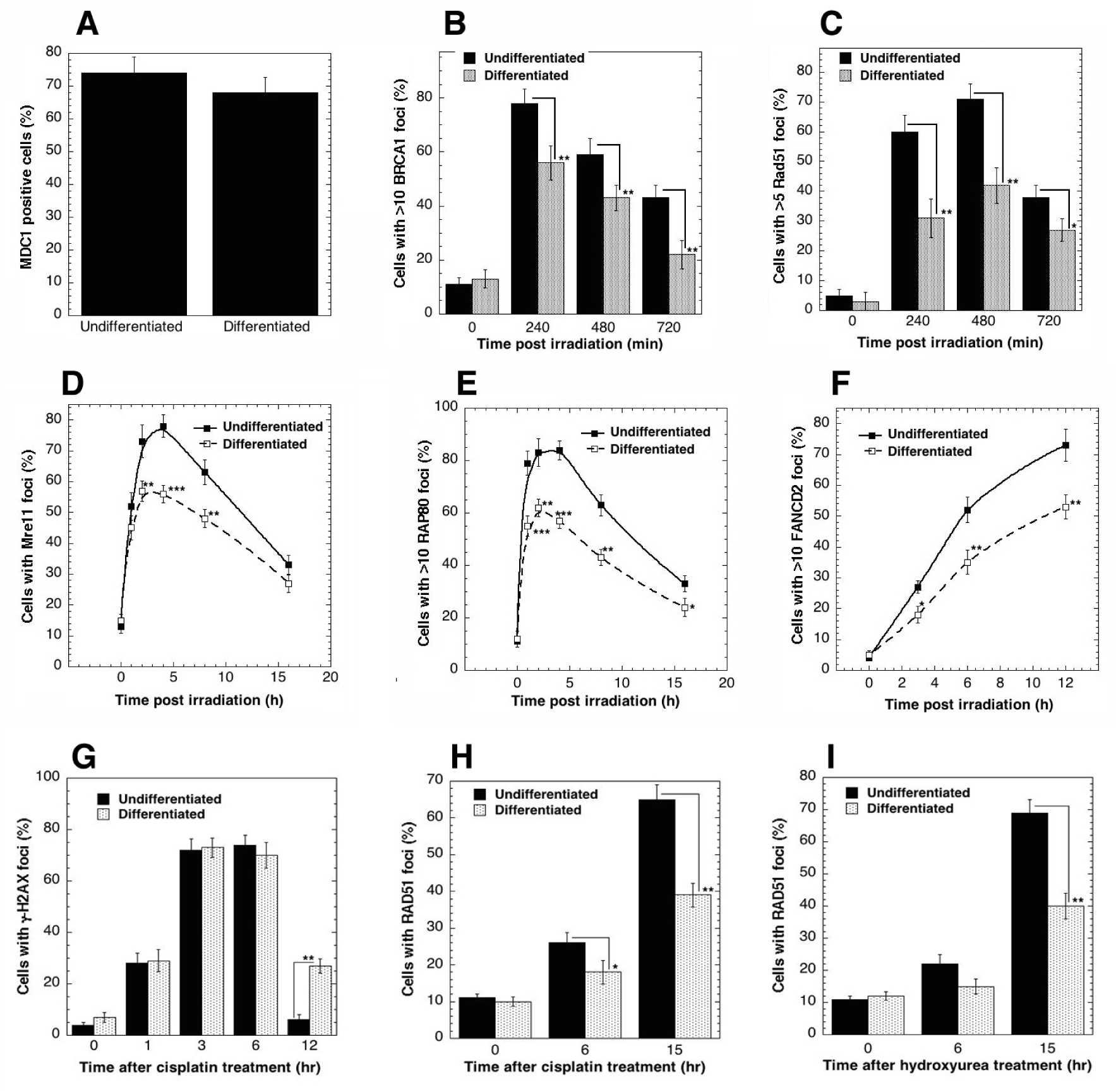
HR repair factor foci formation after DNA damage in undifferentiated and differentiated cells. Stem cells and differentiated cells were exposed to various doses of IR and percentage of positive MDC1 cells **(A)** with BRCA1 foci **(B)**, Rad51 foci **(C)**, Mre11 foci **(D)**, RPA 80 **(E)** and FANCD2 **(F)** stained with respective antibodies; images were captured using a Zeiss Axio Scope fluorescent microscope and scored with the Image G software (v1.47, NIH). B12-2 and differentiated cells were exposed to cisplatin (2 μM) or Hydroxyurea (2 mM) to induce DNA damage and cisplatin induced *γ*-H2AX **(G)** and Rad51 **(H)** and hydroxyurea-induced Rad 51 **(I)** foci signals were measured after staining with respective antibodies. Significance using paired student t-test is shown. *, *P*< 0.05; **, *P* < 0.01, n=3.

### Impact of differentiation on DNA replication fork stalling and resolution

The repair of DSBs generated by resolution of replication forks stalled by DNA damage utilizes the HR pathway (Gupta, Hunt, Hegde, et al. 2014, Hunt et al. 2013). We examined whether stem cell differentiation impacts the repair of DNA intra- or interstrand cross-links (ICLs) or collapsed replication forks due to nucleotide pool depletion. ICLs create obstructions to fundamental DNA processes and are repaired predominantly during S-phase when replication forks converge at ICL sites (Raschle et al. 2008). Cisplatin treatment induced a higher frequency of cells with delayed disappearance of *γ*-H2AX foci in differentiated cells (Fig. 4G). Furthermore, RAD51 foci induction, a marker of HR, after cisplatin or hydroxyurea (HU) treatment was reduced in differentiated cells (Fig. 4 H and I).

To determine whether the defective ICL repair observed in differentiated cells is due to altered restart of stalled replication forks, we measured the frequency of stalled replication forks and new replication origin firing by using the chromatin fiber assay (Henry-Mowatt et al. 2003). Cells were pulse-labeled with 5-chlorodeoxyuridine (CldU) followed by HU treatment for 2 h to deplete the nucleotide pool, and subsequently labeled with 5-iododeoxyuridine (IdU) (Petermann et al. 2010, Singh et al. 2013). Contiguous IdU/CldU signals (Fig. 5 A and B), identifying restarted forks, were significantly lower in differentiated than in undifferentiated ES (H9) or iPS (B12-2) cells (Fig. 5C). Further analysis of the DNA fibers indicated the percentage of stalled forks in differentiated cells after 2 h of HU treatment was higher than in undifferentiated cells, suggesting that differentiated cells resolve stalled replication forks less efficiently (Fig. 5 C and D). In addition, differentiated cells had a shorter DNA tract length distribution indicating reduced replication fork speeds (Fig. 5E and F).

**Figure 5:**
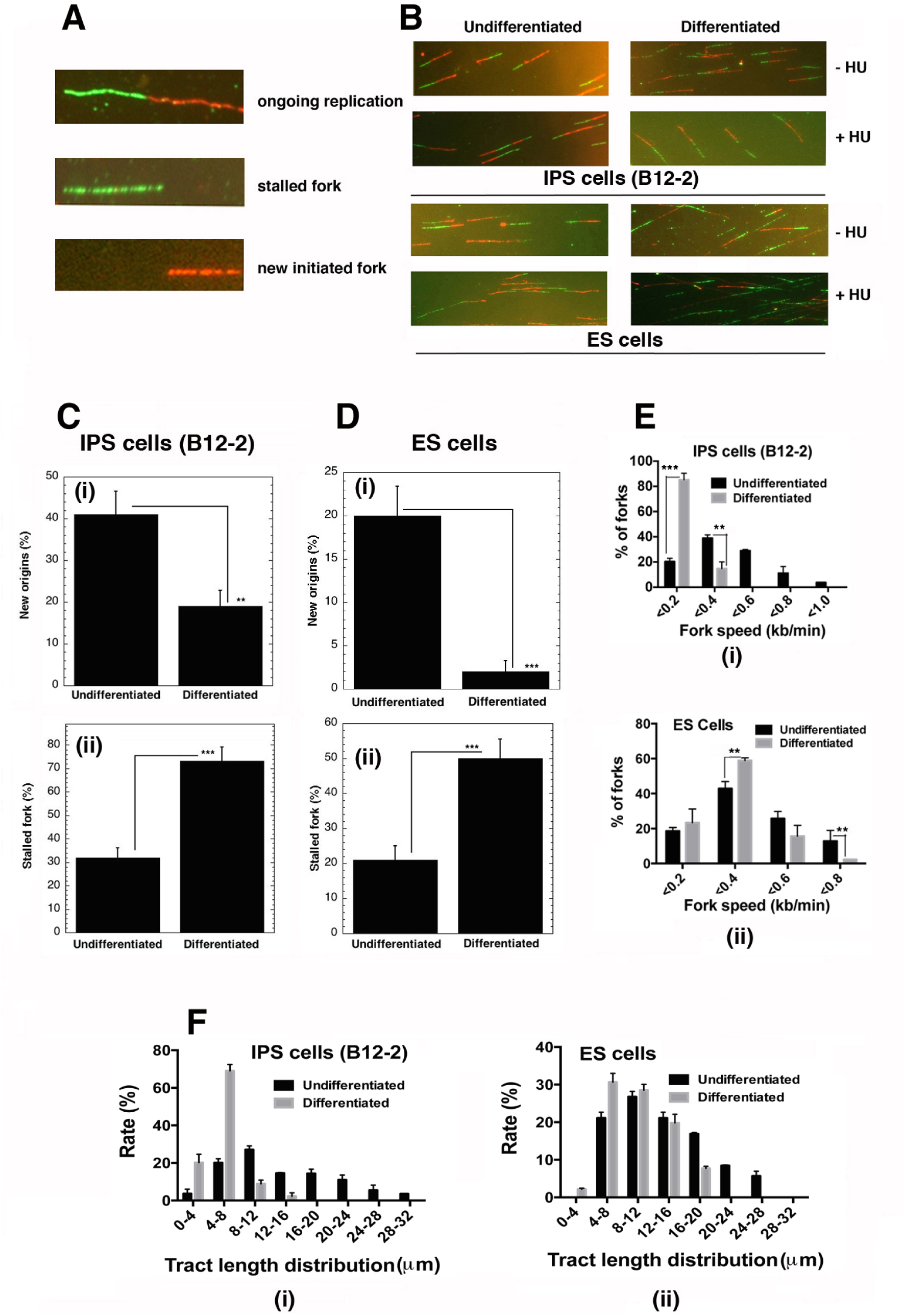
Stalled DNA replication forks and initiation of new origins in B12-2 and differentiated cells: **(A)** DNA labeling and hydroxyurea treatment protocol for single DNA fiber analysis showing ongoing replication (green “IdU” followed by red “CldU” track; green only tracks indicate stalled forks and red only tracks identify newly initiated replication sites); **(B)** representative image of replication tracks from B12-2 (iPSC) and H-9 ES cells and their differentiated progeny in the presence and absence of hydroxyurea, which depletes the nucleotide pool. **(C)** Quantification of new origins as determined by CIdU signal after 2 h of HU treatment in B12-2 and differentiated cells and H-9 and differentiated cells. **(D)** Differentiated cells show a decrease in incorporation of CIdU and maximum frequency of cells with stalled forks. **(E)** Fork speed and **(F)** track length distribution in undifferentiated (ES and iPS cells) and differentiated cells. Significance by paired student t-test is shown. *, *P*< 0.05; **, *P* < 0.01, n=3.

Replication fork stalling can also arise from interstrand crosslinks or RNA: DNA hybrid formation, the latter also referred to as transcriptional R loops. We observed a higher frequency of R-loop formation in differentiated than in undifferentiated cells (Fig. S5).

## Discussion

Ectopic expression of Oct4, Sox2, Klf4 and c-Myc and/or selected groups of transcription factors in somatic cells of mouse or human origin can generate iPS cells (Huangfu et al. 2008, Hanna, Saha, and Jaenisch 2010, Yamanaka 2007, 2012). The iPS clones B12-2 and B12-3 used here were induced to pluripotent state by introduction of two transcription factors (OCT4, SOX-2) and valproic acid (VPA), a small molecule histone deacetylase inhibitor (HDAC) (Huangfu et al. 2008). We show that after differentiation, cells have lower levels of H4K16ac and a correspondingly reduced HR capability.

Both ES and iPS cells undergo epigenetic changes marked by H3K4me1, me2, me3, H3K27me3, HeK36me3 during reprogramming (Koche et al. 2011). As stem cells differentiate, chromatin changes are obvious in conjunction with transcription regulatory changes (Nashun, Hill, and Hajkova 2015, Tran et al. 2015), which can also alter the DDR. This can result in DNA damage accumulation and replicative arrest as observed in Hutchinson-Gilford progeria syndrome (Musich and Zou 2011). Maintenance of genomic stability is of vital importance for stem cells, as mutations can compromise derived cell lineages and their progenitors (Adams et al. 2010, Serrano et al. 2011). DNA damage has been reported to specifically accumulate in stem cell compartments with age (Mandal, Blanpain, and Rossi 2011, Behrens et al. 2014). We report that increased damage could be due to down regulation of DSB repair by HR. We compared the DNA damage response in stem cells before and after differentiation and found that differentiated stem cells have a: (1) higher frequency of spontaneous chromosome aberrations; (2) reduced level of DNA DSB repair after IR exposure; (3) higher frequency of S-phase specific IR-induced chromosome aberrations; (4) higher frequency of residual *γ*-H2AX foci formation after IR exposure or cisplatin treatment; (5) higher frequency of cells with 53BP1 and RIF1 co-localization; and a (6) higher frequency of cells with reduced number of Rad51 or BRCA1 foci after IR exposure or cisplatin treatment as compared to undifferentiated stem cells. The higher frequency of chromosome aberrations found in differentiated cells correlated with reduced DSB repair and a higher frequency of S-phase specific aberrations, suggesting that differentiation impacts DSB repair. The higher frequency of chromosome aberrations is not due to an altered cell cycle distribution as there is no difference in the distribution of cell cycle phases between undifferentiated and differentiated cells. Since no difference in IR-induced G1- or G2-specific chromosome aberrations was observed between undifferentiated and differentiated cells, this suggests the non-homologous end joining DSB repair pathway is not affected, as it is the dominant mode of DSB repair during G-1 or G-2 phase cells. In contrast, differentiated cells have a higher frequency of S-phase specific IR-induced chromosome aberrations, suggesting that differentiation impairs homologous recombination DSB repair, which primarily occurs in S-phase cells. Thus our results suggest that while the NHEJ pathway is minimally altered, DSB repair by HR is impaired by differentiation of stem cells.

Only a subset of differentiated cells had a higher frequency of residual IR-induced *γ*-H2AX foci suggesting that most of the cells can repair the DNA damage. Further, since differentiated cells showed a higher frequency of cells with co-localization of 53BP1 with RIF1, this suggests that in such cells the subsequent recruitment of the HR related proteins is impaired. Consistent with these observations, differentiated cells have a reduced frequency of foci formation by Rad51, BRCA1 and other HR related protein.

Defects in HR can also increase stalled replication forks and we found a higher frequency of stalled replication forks and lower frequency of new replication origins in differentiated stem cells. The increased level of stalled forks could be due to reduced resolution and repair of the forks by HR. Alternatively, increased interstrand crosslinks or transcriptional RNA: DNA hybrids, also called R loops, may contribute to stalling. This latter mechanism seems more likely since differentiated cells exhibited a higher frequency of R-loops. Further, the reduced levels of H4K16ac may be important due to its unique ability to control chromatin structure and protein interactions which may facilitate a more open, repair conducive chromatin configuration (Pandita 2013, Horikoshi, Hunt, and Pandita 2016). Acetylation of H4K16 also limits 53BP1 association with damaged chromatin to promote repair by the HR pathway (Tang et al. 2013). Thus, the reduced levels of H4K16ac in differentiated cells could be one of the factors contributing to aberrant DSB repair.

Fully developed tumors are speculated to emerge only in progenitor populations that differentiate from stem cells. Although this is poorly understood, it may be the result of very tight control mechanisms regulating proper homeostasis in SC compartments and which may be absent in downstream populations. This becomes evident when differentiated cells are exposed to DNA damaging agents as a higher frequency of cells with residual damage is observed. It is also possible that genomic instability results in mutational events, tumorigenesis would then arise during the downstream expansion of progenitors committed to a specific lineage (Lopez-Bertoni, Li, and Laterra 2015). Since tissue is largely constituted of differentiated cells, it is important to develop strategies for avoiding the effects of poor DNA damage repair arising during differentiation in order to maximize the therapeutic potential of stem cells.

## Materials and Methods

### Reagents and antibodies

Antibodies were from Cell Signaling: Chk1 (#2360), pChk1 (Ser317) (#12302S), Chk2 (#2662S), pChk2 (Thr68) (#2661), ATR (#2790S), p-ATR (#Ser428), pATM-Ser-1981 (#13050S), total H2AX (#7631), H2AX-Ser 139 (#9718), OCT 4 (#2890) and nanog (cat# 4903). Santa Cruz antibodies: 53BP1(#sc-22760) ATM (#sc-7230), BRCA1 (#sc-642). Abcam antibodies: RPA-70 (#ab79398), Rad51 (#ab63801). Genetex antibodies: MRE11 (#GTX70212). Bethyl Laboratories antibodies: RIF1 (A300-5671). Upstate Biotechnology and Millipore: H2AX-Ser 139 (# 05-636).

### Cell culture and differentiation of human induced pluripotent (iPSC) cells

Human induced pluripotent cells (clones B12-2 and B12-3) were a kind gift from Dr. Douglas Milton and Dr. D. Huangfu of the Harvard Stem Cell and Regenerative Center, MA. iPS cells (B12-2 or B12-3) were initially grown in 90% knockout DMEM, 10% knock-out serum replacer, 10% Plasmanate (human plasma), 1 mM L-glutamine, 1 mM non-essential amino acids, 0.1 mM β-mercaptoethanol (55 mM Stock), supplemented with 10 ng/ml bFGF on mitotically-inactivated mouse embryonic fibroblast feeder layers. Cells were grown on Matrigel in mTeSR-1 medium (Stem Cell Technologies) for maintenance of iPSC in feeder free culture. The iPS cells grown on Matrigel were dissociated using 2 mg/ml of collagenase IV (Invitrogen), washed and cultured in suspension in ultra-low attachment plates (Corning) in the differentiation medium containing 80% knockout DMEM, 1 mM L-glutamine, 0.1 mM β-mercaptoethanol, 1 mM non-essential amino acids and 20% defined FBS (Hyclone). The media was changed on days 2 and 4 and on day 6 the embryoid bodies (EBs) were transferred onto gelatin coated plates (3-4 EBs per cm^2^) and cultured for additional days as described in the Results Section.

Clone B12-2 and B12-3 were induced by reprogramming primary human fibroblast to pluripotent state (Huangfu et al. 2008). Cultures of undifferentiated ES or iPS cells were designated as day 0 control.

### Western blot analysis

Cell cultures were washed with cold PBS and were lysed in the cell extraction buffer (Invitrogen) supplemented with 1 mM phenylmethyl sulfonyl fluoride and protease inhibitor cocktail (Sigma-Aldrich) for 30 min on ice. Equal protein aliquots were resolved on SDS-PAGE and transferred to nitrocellulose membrane. The membranes were blocked for 45 min in a blocking buffer (5% non-fat dry milk in Tris buffered saline), washed and incubated with specific antibodies for 2 h at room temperature. Proteins were detected with HRP-conjugated secondary antibodies and visualized by enhanced chemiluminescence.

### Detection of γ-H2AX foci and other DDR components

Stem cells and differentiated cells were exposed to 2 Gy-10 Gy (depending upon marker detection) of radiation in chamber slides (Nunc^®^ Lab-Tek^®^ II) and cells allowed to recover for 0.5-12 h at 37° C. Following fixation and permeabilization, cells were probed with antibodies against phosphorylated H2AX-Ser136, RAD 51, 53BP1, RIF1, BRCA1, MDC1, Mre 11, FANCD2 and RPA 80. *γ*-H2AX foci and foci from other proteins were visualized using a Zeiss Axio Scope fluorescent microscope and scored with the Image G software (v1.47, NIH). At least 100 cells were evaluated for each sample to ensure statistical reliability.

### DNA fiber assay

DNA fiber spreads of ES cells (H-9); iPSCs (B12-2) and derived differentiated cells were prepared as described (Singh et al. 2013) with minor modifications. Briefly, ongoing replication sites in cells were labeled with IdU (50 μM) followed by exposure to hydroxyurea (4 mM), washing and labeling with CldU (50 μM). Fibers were quantified using Image J software.

### Comet assay

The alkaline Comet assay measures DNA strand breaks in single cells (Comet Assay Kit, Trevigen). Comet tail moments were measured in at least 100 cells and quantified using the CometScore software.

### Chromosomal aberration analysis at metaphase

Analysis was performed as described previously (Pandita et al. 2006) (Singh et al. 2013). Cells were irradiated at the dose of 3 Gy and analyzed for metaphase aberrations after 12 h (Gupta, Hunt, Chakraborty, et al. 2014). Cisplatin-induced chromosome aberrations were analyzed as described (Singh et al. 2013).

### Isolation of glial cells from mouse brain

Mixed cortical cell isolation culture of astrocytes was performed using P3-P4 mouse pups. Pups were sacrificed by decapitation and the heads were sprayed with 70% ethanol. Cortices were dissected and placed in the petri dish containing DMEM media (10% FBS, 1% Penn Strep). Cortices were chopped into a homogenous paste with the help of a fine blade and the paste was transferred into a 15 ml conical tube. The cells were dissociated with trypsin (0.25% trypsin-EDTA) for 5 min and neutralized with the addition of fresh medium. Finally, cells were plated on the petri dishes and medium was changed after 24 h to remove cell debris. After one week, when the cells became confluent, they were trypsinized and plated on a 24-well plate with circular coverslip for immunostaining. Cells were irradiated with 2 Gy of IR and fixed at different time intervals (0-24 h) for immunostaining of phosphorylated γ-H2AX and Rad 51.

## Acknowledgement

This work was supported by funds from The Houston Methodist Research Institute and grants (CA129537 and GM109768) from National Institutes of Health.

## Supplementary Figure Legends

**S1:**
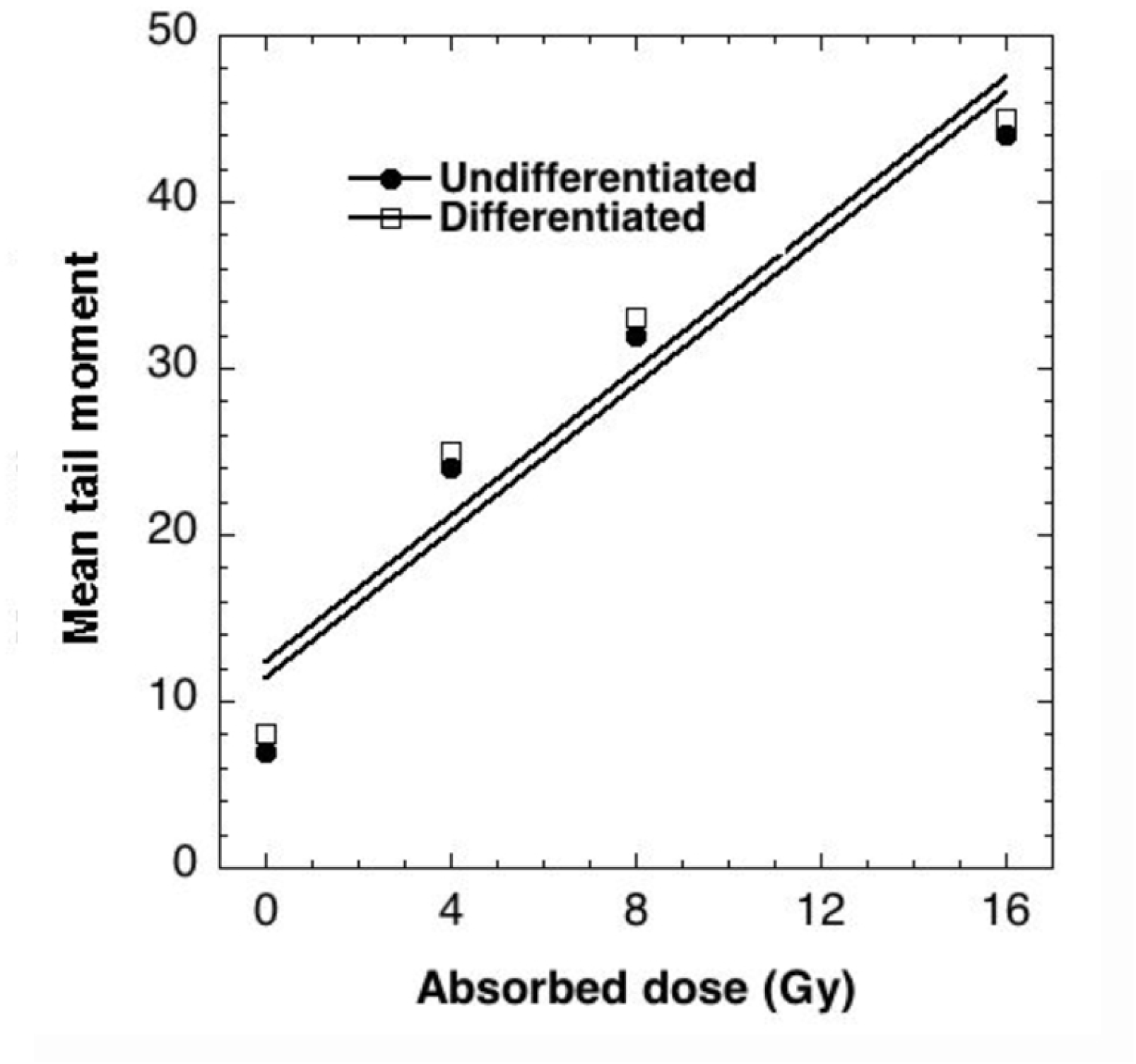
Absorbed dose of radiation in stem and differentiated cells.

**S2:**
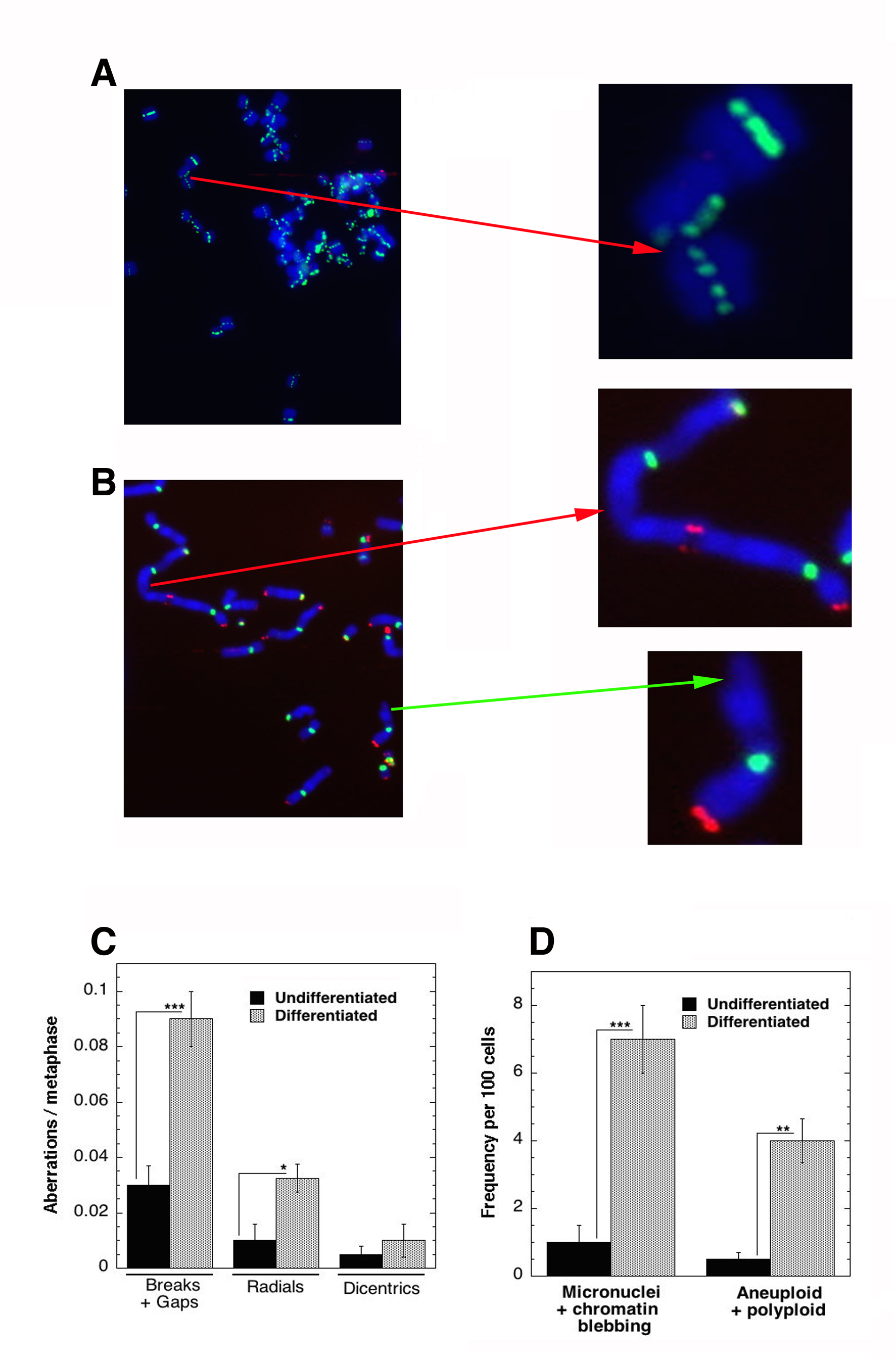
Polyploidy and telomere loss in response to induced stress in differentiated cells and comparison of genomic stability in undifferentiated and differentiated stem cells.

**S3:**
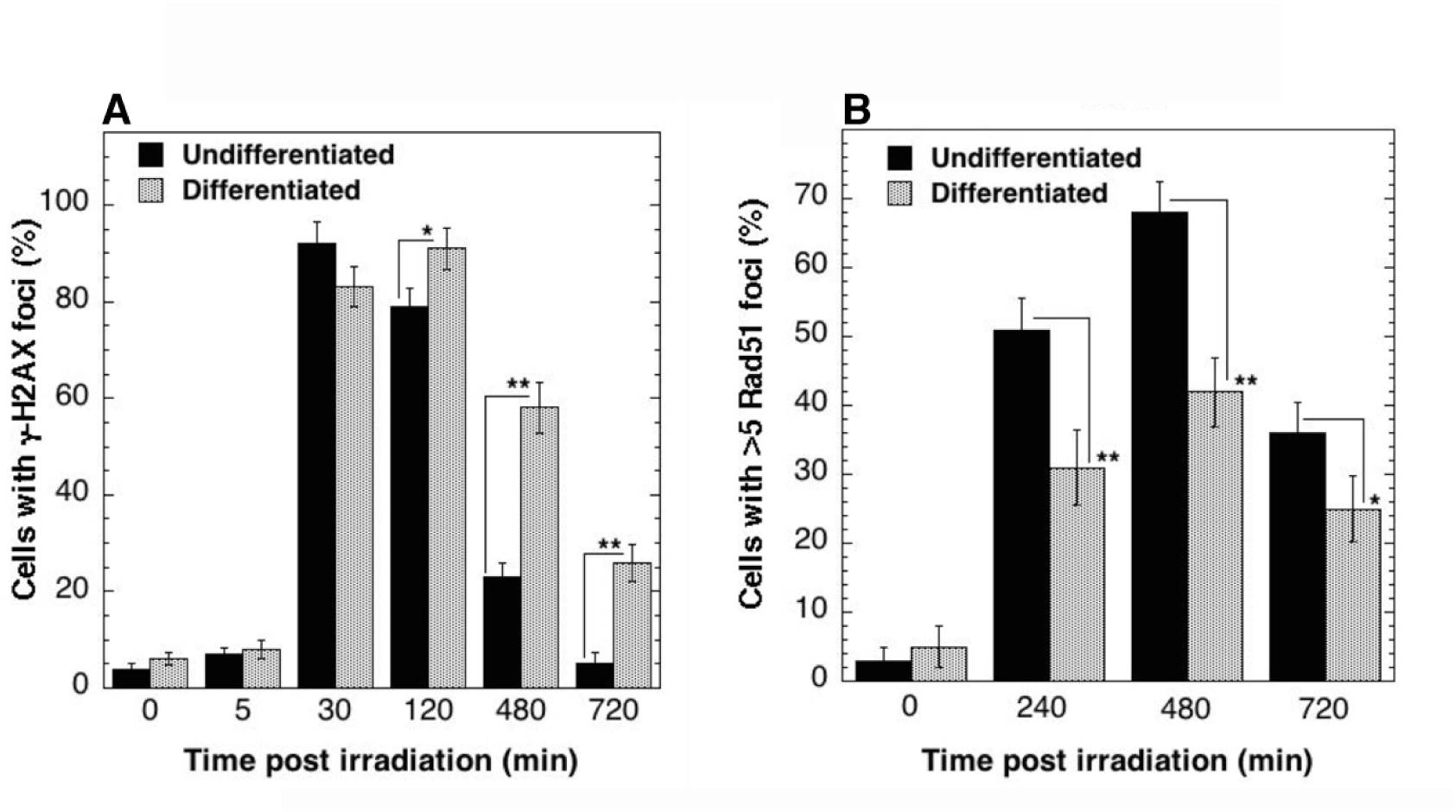
DDR in undifferentiated and differentiated ES cells.

**S4:**
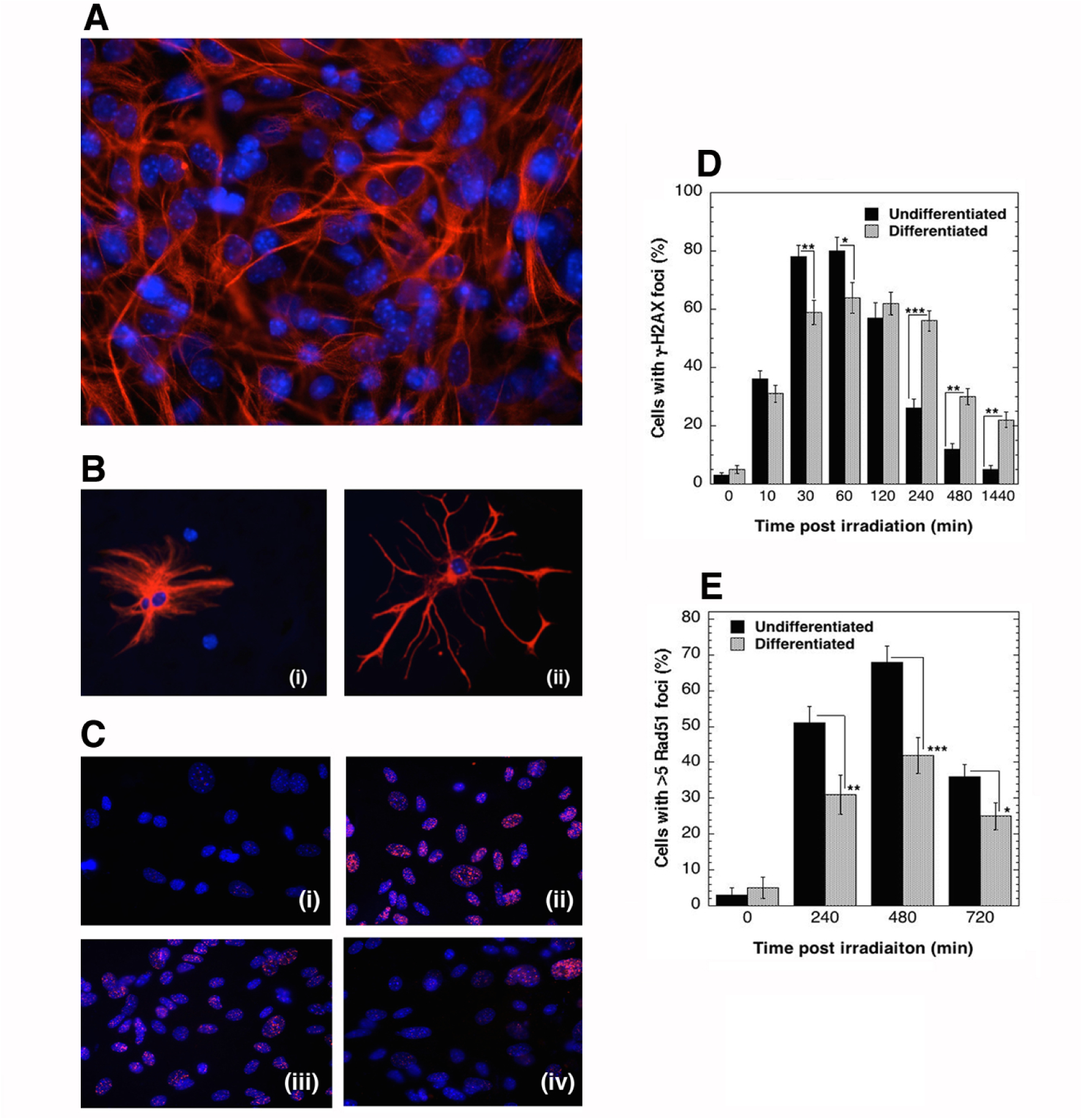
DNA damage response in early (day 7) and late (day 30) mouse astrocytes: (A) Astrocyte culture from mouse p3-p4 pups stained with glial fibrillary acidic protein (GFAP) (B) (i) astrocyte initial stage and (ii) later stage. (C) Representative images of γ-H2AX foci (0-720 min post irradiation); (D) & (E) early stage (undifferentiated) and late stage (differentiated) astrocytes exhibiting γ-H2AX foci (D) and Rad 51 (E) foci 0-12 h post irradiation.

**S5:**
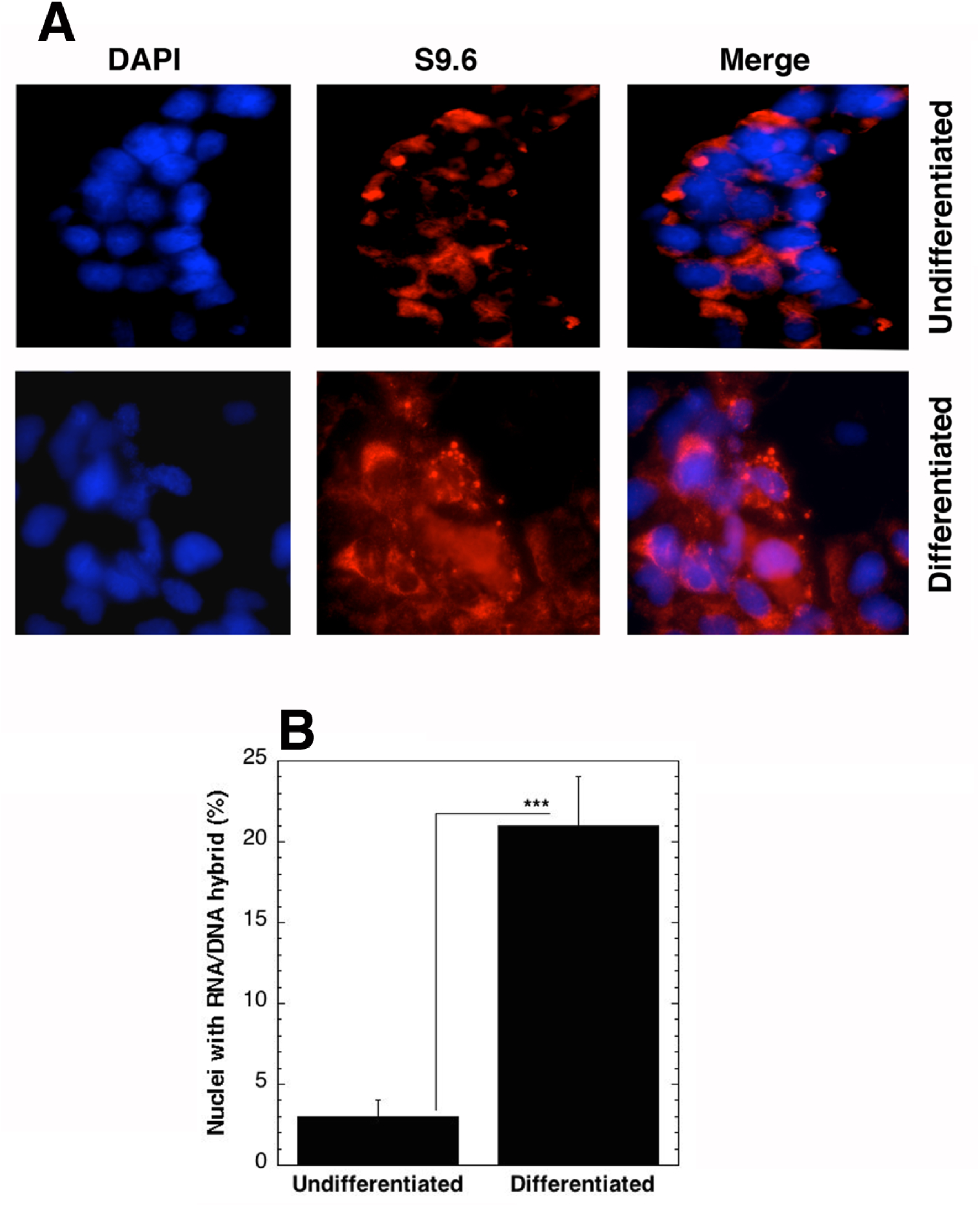
R- loop formation (RNA/DNA hybrids) in undifferentiated and differentiated iPS cells.

